# Characterization of a water-soluble formulation of *Passiflora incarnata* and its effect on sperm motility: A novel male-centric contraceptive formulation

**DOI:** 10.1101/2025.11.23.687804

**Authors:** Srinjoy Chakraborty, Harsh Kumar Singh, Kalpana Nagpal, Sudipta Saha

## Abstract

*Passiflora incarnata* (Passionflower) has a long history of use in traditional medicine across South America and Europe, primarily for its sedative and hypnotic effects. Recent evidence suggests that its bioactive constituents can modulate cellular signaling pathways. However, its impact on reproductive physiology, particularly sperm motility, remains underexplored. This study aimed to investigate the effects of an aqueous *P. incarnata* formulation on sperm motility and viability, with the goal of evaluating its potential as a plant-based, non-hormonal contraceptive agent. An aqueous extract of *P. incarnata* was prepared and subjected to phytochemical analysis to quantify polyphenols, flavonoids, and carbohydrate content using standard assays. Antioxidant activity was assessed via free radical scavenging assays. The presence of bioactive macromolecules was confirmed by liquid chromatography–mass spectrometry (LC-MS). Goat spermatozoa, isolated from the caudal epididymis, were treated with the formulation. Sperm motility (horizontal and vertical) was evaluated microscopically and spectrophotometrically, respectively. Sperm viability was assessed using standard viability staining. Phytochemical screening revealed high polyphenol (16.85 mg gallic acid equivalent/g) and flavonoid (2.5 mg quercetin equivalent/g) content, along with notable antioxidant activity (87.25%). LC-MS confirmed the presence of diverse plant macromolecules. Treatment with the *P. incarnata* formulation significantly reduced both horizontal (28.5%) and vertical (20%) sperm motility (both p < 0.0001), while sperm viability remained unaffected. The water solubility of the formulation supports its development as a topical contraceptive gel. The aqueous *P. incarnata* formulation modulates sperm motility without compromising cell viability, likely via biochemical pathway regulation. These findings highlight its potential as a novel, non-hormonal, plant-based contraceptive.

## 1. Introduction

The use of plant metabolites for altering cellular physiology or sperm motility to control fertilization has been studied extensively in the past few decades. The use of contraceptives is one of the strategies to control pregnancies. Importantly, the extraction and application of plant macromolecules opens extensive avenues to control animal cell behavior. This study focuses on utilizing plant-derived macromolecules to alter sperm cell motility by specifically controlling the sperm flagellar motility.

Sperm cells are highly specialized, haploid, terminally differentiated and mostly transcriptionally inactive cells. The key functions performed by the sperm cells, ranging from capacitation, acrosome reaction and eventually fertilization usually occur outside the human male body. One of the key factors that determines successful fertilization is sperm motility. It is known that immotile sperm cells cannot travel through the female uterotubal tract, thereby rendering them ineffective. The sperm flagellum plays a critical role in regulating motility. The complex mechanisms that contribute to sperm flagellar movement have been studied extensively. However, these molecular mechanisms can be exploited using various ligands and molecules to control sperm motility by either enhancing or reducing motility [1].

Male contraceptives are a class of chemical products the render a sperm cell immotile or hamper the production of sperm cells making them unable to fertilize an ovum [2]. Synthetic spermicides that have been commercially available, such as nonyxenol-9 (N-9) have been reported to cause problems such as vaginal irritation and inflammation. These drugs can also disrupt the vaginal epithelium and may cause detrimental effect to the normal vaginal flora [3]. An ideal spermicidal or male-specific contraceptive agent must incapacitate or immobilize sperm cells and should not be harmful to the normal physiology of the vagina or the penile mucosa and should also be commercially inexpensive so that a majority of the population can use it without any detrimental effect.

With the ever-increasing population and lack of family planning awareness among many, the availability of an oral or topical herbal contraceptive would be ideal candidate to control unplanned pregnancies. Most contraception methods involve the use of hormonal interventions or surgery. The effectiveness of basic methods such as the barrier method, perhaps the most widely used, is questionable. Furthermore, most hormonal contraceptives are women-centric and involve the alteration of the menstruation cycle, giving rise to problems such as irregular menstruation cycles, premenstrual syndrome (PMS), dysmenorrhea (menstrual cramps), heavy or long periods, anemia (low hemoglobin), premenstrual dysmorphic disorder (PMDD), endometriosis, uterine fibroids, acne, migraines, unwanted hair growth, menopause-related hot flashes, and /or risk of uterine, ovarian, and colon cancer [1,4,5]. A study in Europe involving 6676 women reported several side effects of conventional oral contraceptive agents such as hirsutism, nausea, bleeding irregularities, breast tenderness mood changes, and weight gain [6]. Several studies have reported the antifertility activity of several plant extracts, including *Carica papaya* seeds, *Balanites roxburghii*, and *Phyllanthus amarus* in animal models. Several traditional texts have various references of using plant products as contraceptives; however, relatively little research has been done to verify their effectiveness and determine their mechanisms of action [7–9].

This study experimentally demonstrates the ability of a water-soluble passion flower formulation as a male-centric contraceptive formulation that can significantly reduce sperm motility. Passionflower extract has been utilized as a traditional remedy for anxiety and sleep disorders. Several studies have highlighted its anxiolytic properties, attributing them to the presence of compounds such as flavonoids, alkaloids, and coumarins. These bioactive metabolites act on the central nervous system, modulating neurotransmitter levels and promoting relaxation without the sedative side effects associated with conventional anxiolytic medications [10,11]. Moreover, *Passiflora incarnata* has demonstrated potential in managing insomnia. The flavonoids present in the plant extract, particularly chrysin, have been shown to bind to benzodiazepine receptors, inducing sedation and improving sleep quality. This natural alternative provides a promising avenue for individuals seeking sleep aids without the risk of dependence or adverse effects commonly associated with synthetic pharmaceuticals. Furthermore, *Passiflora incarnata* has shown promise in the realm of pain management. The plant’s extract possesses analgesic properties, which can be attributed to the presence of alkaloids like harman and harmine. These compounds interact with pain receptors, modulating the perception of pain and providing relief without the adverse effects associated with conventional analgesics. The antispasmodic effects of passionflower extract also make it a valuable candidate for gastrointestinal disorders. Research suggests that *Passiflora incarnata* can alleviate symptoms associated with irritable bowel syndrome (IBS) and other gastrointestinal issues by relaxing smooth muscle contractions in the intestines[11–13].

The exploration of plant extracts as alternatives in controlling sperm motility holds promise for revolutionizing reproductive health. The biochemical basis of their action, involving modulation of ion channels, reduction of oxidative stress, and interaction with membrane receptors, underscores their potential in influencing sperm movement. As the demand for sustainable and personalized reproductive health solutions grows, plant extracts emerge as valuable candidates for further research and development in the field. Their unique biochemical properties position them as a bridge between traditional methods and innovative, nature-inspired approaches to controlling sperm motility [14]. This study is aimed at determining the effect of *Passiflora incarnata* (PI) formulation in regulating sperm motility. This study highlights the role of using naturally available plant extracts that can be used as an alternative contraception method.

## 2. Materials and Methods

All chemicals used in the study were procured from SRL, CDH, and/or sigma chemicals. All chemicals were of AR grade, unless otherwise stated. Commercially available extract of *Passiflora incarnata* in powdered form was obtained from Xi’an Aladdin Bio-Tech Co. Ltd. A 1 mg/mL w/v solution made using distilled water or DMSO was used for each of the experiments performed during the tenure of the study. Adult goats from the local slaughterhouses were utilized for the experiment as they do not require any ethical clearances owing to their utility as a raw food material. Fresh testes and epididymis were obtained. The spermatozoa were extracted within two to four hours of slaughtering the animals.

### 2.1 Solubility of Passiflora incarnata extract

The primary aim of this study was to find a suitable additive for sperm cell storage obtained from natural sources. Therefore, to determine the solubility of the *Passiflora incarnata* powder, multiple solvents were used. Because sperm cells are sensitive to solvents such as DMSO, methanol and other organic solvents, which are routinely used to obtain phytochemicals, a soluble plant product worked in favor of the study. Using the shake flask method, the equilibrium solubility of the *P. incarnata* extract was found in double distilled water, modified Ringer’s phosphate solution (which consists of sodium chloride (119 mM); potassium chloride (5 mM); magnesium sulphate (1.2 mM); glucose (10 mM); potassium phosphate (16.3 mM); penicillin (50 units/mL, pH 7.4); methanol; ethanol; and dimethyl sulfoxide). 5 mg of the solute was added into varying volumes of solvent till a saturation point was reached. The volume showing maximum solubility was noted.

### 2.2. UV-Visible Absorption analysis

A 1 mg/mL solution of *Passiflora incarnata* in modified Ringer’s phosphate solution was subject to UV-Visible Spectroscopic analysis using a *Labman, Chennai, India* single beam UV-Visible spectrophotometer. A wavelength range from 200 nm to 800 nm was used for the study. Ringer’s Phosphate solution was used as blank for the experiment.

### 2.3 Fourier Transform Infrared Spectroscopic Analysis

FT-IR spectra of the dry *Passiflora incarnata* extract were obtained using a Perkin Elmer FTIR-ATR/FRONTIER IR spectrophotometer (Perkin Elmer, Massachusetts, USA). FT-IR spectra was obtained within the range of 500–4000 cm^−1^ at 2 cm^−1^ resolution.

### 2.4 Total Phenol Estimation of P. incarnata formulation

Folin–Ciocalteu technique was used to determine the total phenolic content of solutions containing the *P. incarnata* formulation. A standard graph was prepared using a 1 mg/mL (w/v) solution of gallic acid. This standard graph was used to determine the total phenolic content by using the gallic acid equivalents method. A 10 mg/mL (w/v) crude extract solution of the *P. incarnata* extract was prepared for the study. The entire experiment was performed in dark conditions. Initially, 100 μL of Folin–Ciocalteu (1:1) reagent was added to different concentrations of gallic acid for 3 min. Then, 1 mL of 10% (w/v) sodium carbonate was added to the tubes. The tubes were incubated in the dark for 1 hour. Each sample was tested for its absorbance value at 760 nm using a *Labman, Chennai, India* single beam UV spectrophotometer. The total phenolic content was calculated and expressed as mg of gallic acid equivalent per gram of dry weight with the help of the gallic acid standard curve.

### 2.5 Total Flavonoid Estimation in P. incarnata formulation

The aluminum chloride colorimetric method was used to determine the total flavonoid content in the aqueous formulation. Quercetin was employed to construct the standard calibration curve. A 1 mg/mL (w/v) solution of quercetin in DMSO was prepared as the working stock. To the dilutions, 100 μL of 10% aluminum chloride and 1 M sodium acetate were added. After briefly mixing the contents, deionized distilled water (DDH O, 2.8 mL) was added, and the mixture was incubated at STP conditions for one hour. The reaction mixtures were tested for their absorbance values at 415 nm wavelength using a UV-Vis spectrophotometer. The total flavonoid content of the *P. incarnata* formulation was expressed as milligrams of quercetin equivalents per gram of dry weight.

### 2.6 Qualitative Test to Determine the Presence of Saponin

To generate a stable and persistent froth, *P. incarnata* formulation (1 mg/mL in RPS medium) was subjected to vigorous shaking. The froth so obtained was mixed with three drops of olive oil and vigorously shaken. The presence of saponins formation was indicated by the consistent emulsion appearance.

### 2.7 Antioxidant scavenging assay

1, 1-diphenyl-2-picrylhydrazyl (DPPH) assay: This assay technique was used to determine free radical scavenging activity of the *P. incarnata* formulation solution. The evaluation was performed on the basis of the presence of its hydrogen-donating or radical-scavenging capabilities, utilizing the stable free radical DPPH. 200 µL of the 1 mg/mL *P. incarnata* formulation solution was added to DPPH solution (0.08 mg/mL in ethanol). The reaction mixture was incubated at room temperature for 30 minutes in complete darkness. Following incubation, the tubes were centrifuged at 4000 rpm for 10 minutes. Centrifugation was carried out to obtain 0.5 mL of the supernatant and was transferred into the fresh tubes containing 1 mL of ethanol. The absorbance of the supernatant so obtained was measured at 517 nm wavelength using ascorbic acid as the reference. The percentage inhibition of the DPPH free radical was calculated against ethanol as the blank using the following equation:

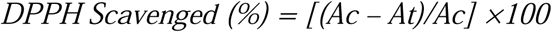

H_2_O_2_ scavenging activity was evaluated by preparing a solution of H_2_O_2_ (40 mM) in phosphate buffered saline (pH 7.4). To this, varying concentrations (16 – 1000 μg/mL) of formulation (3.4 mL) in phosphate buffered saline were added along with H_2_O_2_ (0.6 mL, 40 mM). After a 10-minute incubation period, the absorbance at 230 nm was measured against phosphate buffered saline without hydrogen peroxide. The following formula was used to calculate the percentage of H_2_O_2_ scavenging by the *P. incarnata* formulation as well as the standard ascorbic acid:

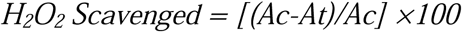

### 2.8 Fehling’s Test for testing the presence of carbohydrates

Fehling’s test is an assay technique which is used to distinguish between reducing sugars from non-reducing sugars; as well as to differentiate between carbohydrates possessing ketone functional groups versus those with water-soluble functional groups. Fehling’s reagent A (CuSO4·7H_2_O (7 g) in 100 mL of water) and Fehling’s reagent B (KOH (24 g) & potassium sodium tartrate (34.6 g) in 100 mL of water) were prepared and were mixed in equal volumes to form the complete Fehling’s reagent. Then, 1 mL of 5% *P. incarnata* solution was pipetted into a clean and dry test tube, followed by the addition of 2–3 drops of Fehling’s solution. The test tubes were then placed in a boiling water bath for 1–2 minutes, and any change in the color of the solution was observed and recorded.

### 2.9 Protein estimation by Lowry’s method

The protein content of the *P. incarnata* formulation was determined using Lowry’s method for protein quantification. Preparation of the experimental solutions proceeded as follows: Reagent A was formulated by the addition of Na_2_CO_3_ (2%) in NaOH (0.1 N). Reagent B was prepared by addition of CuSO_4_·5H_2_O (0.5%) in sodium citrate solution (1.0%). Reagent C was prepared by the addition of 1 mL of Reagent B to 49 mL of Reagent A. Reagent D was formulated by diluting Folin-Phenol Reagent (2 N Solution, Folin/Ciocalteau) with an equal volume of DDH_2_O.

### 2.10 Identification of phytochemical constituents using LC-MS

Liquid Chromatography was performed using a hypersil gold C18 column, P.S:1.9u, Diameter(mm):100X2.1. (Ultimate 3000 -Thermo Scientific; LTQ Orbitrap XL -Thermo Scientific) Results were then analyzed using Xcalibur 2.1

### 2.11 Isolation of sperm cells from Goat testes

Spermatozoa were collected from the cauda epididymis of goats following the method described by Saha et al., 2013. The procedure involved isolating highly motile spermatozoa from the epididymides at room temperature (32°C ± 1°C) using a modified Ringer’s solution. The spermatozoa counts were determined using a hemocytometer. The concentrations of epididymal plasma used in the assays were quantified based on their protein content, which was assessed using Lowry’s method [15].

### 2.12 Collection of Epididymal Plasma and spermatozoa

To mitigate potential artifacts arising from the adhesion of sperm to the glass surfaces, assay for motility of sperm was conducted in the presence of 6 mg protein per 100 mL epididymal plasma extracted from the caudal epididymis. This plasma was found to possess effective anti-adhesive properties, leading to nearly complete inhibition (approximately 100%) of sperm adhesion to the glass surface of the hemocytometer, as detailed previously [16]. The caudal epididymis was excised and placed in a Petri dish containing Ringer’s phosphate buffer (pH 7.4) at room temperature. The tissue was minced using surgical scissors to release mature sperm cells and epididymal plasma into the buffer solution. The resulting cloudy suspension was filtered through cheese cloth to remove tissue debris. Centrifugation of the filtrate was then carried out (10,000 RPM, 20 minutes, 4°C). The clear supernatants, which contained the epididymal fluids, were collected and stored at –20°C until further experimentation.

### 2.13 Motility-inhibiting activity of P. incarnata formulation by microscopic method

The study assessed the motility-inhibiting effects of *P. incarnata* formulation by examining forward and total motility of spermatozoa using a hemocytometer as the counting chamber. Spermatozoa (0.5 million cells) were exposed to EP in the presence or absence of specified concentrations of the test samples (1 mg/mL aqueous solution of *P. incarnata* formulation) at 32°C ± 1°C for 60 seconds in a total volume of 0.5 mL of the ringer’s physiological solution. Sperm cells exhibiting various motility patterns (total motility) and forward motility were enumerated at 400 times magnification under the phase-contrast microscope. Subsequently, the percentage of motile sperm was calculated based on the observed counts.

### 2.14 Sperm Vertical Velocity Measurement

The study assessed the impact of *P. incarnata* formulation on sperm motility by employing a spectrophotometric technique as detailed previously [17]. A wavelength of 545 nm, corresponding to the peak absorbance of the sperm sample, was selected using the spectrophotometer. A saturation curve depicting absorbance vs time was generated for a scanning period of 3-20 minutes. The modified RPS (1.5 mL) was taken in a cuvette and was scanned spectrophotometer’s cuvette holder to ensure that the light beam passed through the uppermost layer of the solution under normal conditions. After this, 50 µL of spermatozoa (from a stock cell density of 200 × 10^6 cells/mL), suspended in RPS medium and supplemented with Ficoll-400 (2%), was slowly layered at the bottom of the cuvette using a 500 µL Hamilton syringe, both with and without the test sample. Experimental data in terms of absorbance versus time were recorded during each scanning cycle.

### 2.15 Assessment of Sperm Viability

The Trypan blue dye exclusion method was used to check the viability of sperm cells post *P. incarnata* treatment. Briefly, equal volume of sperm suspension (100 uL) was mixed with 0.4% Trypan blue stain and incubated at room temperature for 4 minutes. This mixture was then centrifuged at 600g for 3–4 min. After discarding about 9/10^th^ of the supernatant, a smear was prepared from the pellet to achieve a higher number of sperm cells per field. The smear was then dried in a hot air oven (80, 5 minutes). Sperm cells were then counted (about 200 sperms counted per slide) using a hemocytometer under a bright field microscope at 40× magnification.

### 2.16 Statistical Analysis

All the experiments representing biological triplicates and the observations were expressed as mean ± SEM. Paired t-test and one-way ANOVA was performed to determine the significance level in terms of *P* value using the online Graphpad Prism Software.

## 3. Results

### 3.1 *Passiflora incarnata* formulation was found to be soluble in water and Ringer’s phosphate-buffered saline

To avoid the ill effects of extraction solvents that are regularly used to extract plant metabolites, we used the commercially available powdered form of the *Passiflora incarnata*. The *P. incarnata* extract showed high solubility (Up to 5 mg/mL) in water and modified Ringer’s phosphate solution (RPS). The powder was also soluble in methanol (5 mg/mL), ethanol (5 mg/mL) and DMSO (10 mg/mL). This is especially important because upon scale-up this can be used in a water soluble form for better use.

### 3.2 UV/visible spectrophotometer and Fourier Transform Infrared spectroscopy indicated the presence of various plant metabolites of pharmaceutical importance

The *P. incarnata* formulation was then subject to UV/visible and Fourier Transform Infrared spectroscopy. The UV/visible spectrophotometric analysis revealed a very distinct peak at 280 nm and two insignificant peaks at 320 nm and 327 nm. The peak region corresponding to 280 nm suggests the presence of polyphenols, flavanols, anthocyanin compounds, and phenolic acids in the *P. incarnata* formulation. FT-IR spectra obtained by the KBr pellet indicate the presence of the following functional groups hydroxyl O-H, amine N-H, alkyl double bond C=C, and fluoro substituted alkyl C-F group (**Fig 1A**). The Fourier Transform Infrared (FTIR) spectrum reveals distinct absorption bands characteristic of organic functional groups, enabling identification of molecular bonds and structural motifs. The spectrum spans the mid-infrared region (4000–500 cm□^1^), with prominent peaks indicative of O-H, C-H, C=C, and C-O/C-N vibrations. These observations align with standard FTIR absorption patterns for aromatic and oxygen/nitrogen-containing compounds. A broad, intense absorption band at 3248 cm□^1^ corresponds to O-H stretching vibrations in hydrogen-bonded hydroxyl groups, typically observed in phenolic compounds or alcohols. The region between 3000–2800 cm□^1^ exhibits two distinct absorption patterns: Aromatic C-H stretches (>3000 cm□^1^), observed as weak-to-medium intensity peaks, correspond to sp^2^ hybridized carbons in benzene rings. Aliphatic C-H stretches (<3000 cm□^1^), appearing as sharper peaks near 2920 cm□^1^ and 2850 cm□^1^, arise from sp³ hybridized carbons in methyl (-CH□) or methylene (-CH□-) groups. These aliphatic signals suggest the presence of alkyl side chains or ether linkages, as observed in diphenhydramine and cyclizine. The sharp absorption at 1085 cm□^1^ corresponds to C-O-C asymmetric stretching in ether linkages, a hallmark of 4-phenoxyphenol and diphenhydramine. The complex absorption pattern below 900 cm□^1^ serves as a molecular “fingerprint,” with key peaks at 825 cm□^1^ (para-substituted benzene rings) and 750 cm□^1^ (aromatic C-H out-of-plane bending). These bands confirm the presence of aromatic systems with specific substitution patterns, consistent with the diaryl ether structure of 4-phenoxyphenol. While these features align most closely with 4-phenoxyphenol, spectral overlaps in the C-O/C-N region prevent definitive exclusion of diphenhydramine or cyclizine. For example, cyclizine’s piperazine ring could contribute to the 1240 cm□^1^ peak, while diphenhydramine’s ether and tertiary amine groups might explain the 1085 cm□^1^ and 1240 cm□^1^ bands. The liquid chromatography-mass spectrometry (LC-MS) data reveals critical insights into the molecular composition of the analyzed sample. The total ion chromatogram (TIC) exhibits a primary peak at 11.39 min (relative abundance: 100%), accompanied by several minor peaks across the 14-minute runtime (Fig. 1A). The full-scan mass spectrum (Fig. 1B) displays a base peak at m/z 167.01 and multiple adduct/fragment ions, suggesting the presence of a dominant compound with a molecular weight of approximately 166 Da, indicating the presence of 4-phenoxyphenol. The base peak at m/z 167.01 aligns with the characteristic fragment ion of cyclizine ([C□□H□□N□]□), which undergoes collision-induced dissociation (CID) to lose its methylpiperazine moiety (-C□H□□N□), yielding a stable benzhydryl fragment at m/z 167. High-resolution MS data from reference spectra confirms cyclizine’s protonated molecular ion [M+H] at m/z 267.19, consistent with its molecular formula (C□□H□□N□). The observed sodium adduct at m/z 301.14 further supports cyclizine’s presence, as piperazine derivatives readily form metal adducts under electrospray ionization (ESI) [18]. Diphenhydramine ([M+H]□ = m/z 256.0) shares a fragment ion at m/z 167.0, corresponding to cleavage of its ether bond and subsequent loss of the dimethylaminoethyl group. While the base peak at m/z 167.01 could theoretically arise from diphenhydramine, the absence of a prominent precursor ion at m/z 256 in the full-scan spectrum weakens this hypothesis. The minor peak at m/z 279.16 may represent a sodium adduct ([M+Na] = 256 + 23 = 279), but this aligns ambiguously with diphenhydramine’s expected adduct profile. In conclusion, LC-MS Structure analysis indicated the presence of 4-phenoxyphenol, Diphenhydramine and Cyclizine compounds (**Fig 1B**).

**Fig 1:**
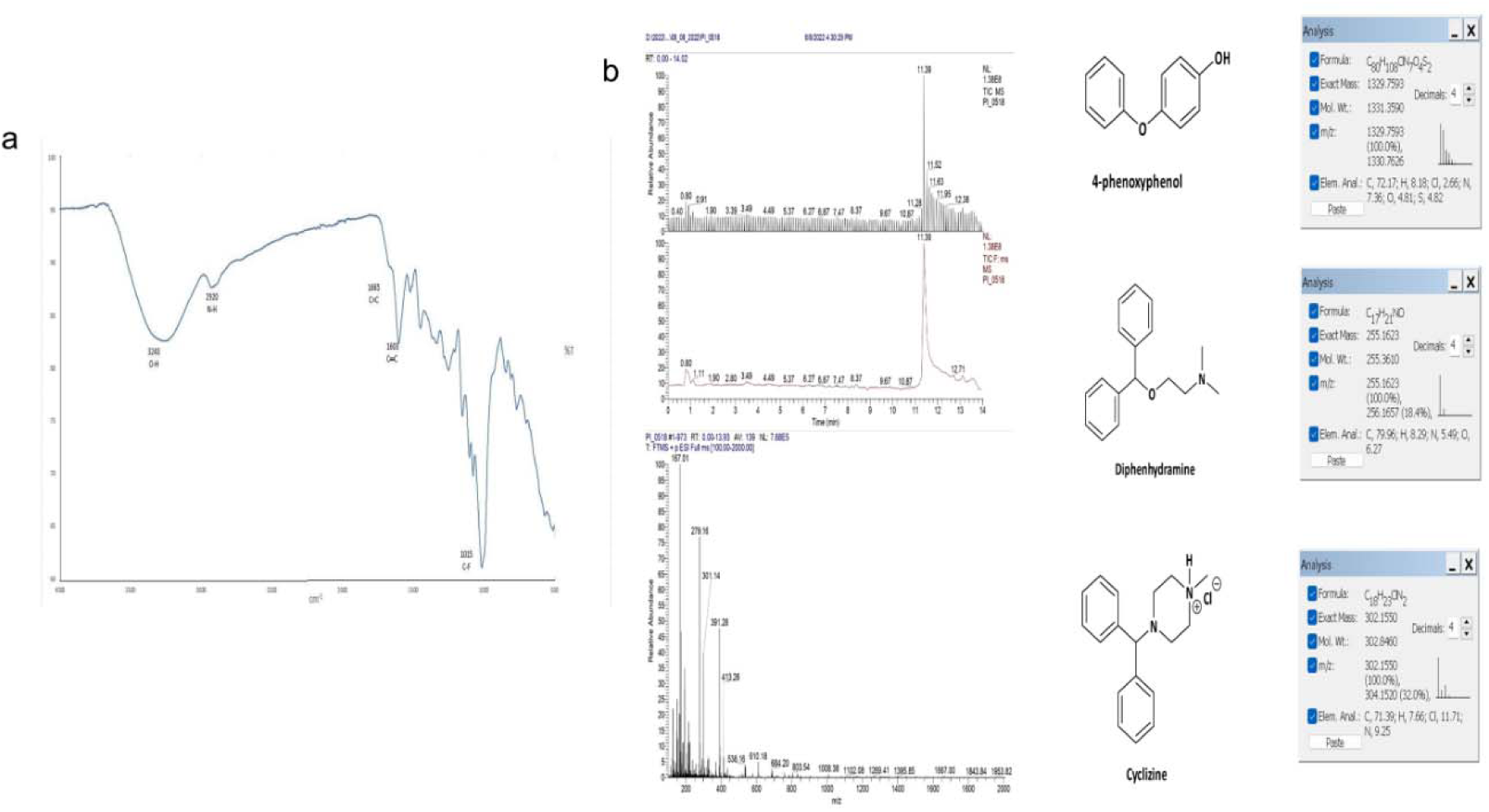
(A) FTIR spectra of *Passiflora incarnata* indicated the presence of O-H, N-H, C=C, C=C, and C-F groups. (B) LC-MS analysis of *Passiflora incarnata* extract indicated the presence of 4-phenoxyphenol, Diphenhydramine and Cyclizine compounds.

### 3.3 Biochemical characterization of *P. incarnata* formulation showed significant levels of phenol, flavonoid, carbohydrates and saponin

The observed value of the total phenolic content of *P. incarnata* formulation was 16.85 mg Gallic acid equivalent /g of dry weight. Similarly, the total flavonoid content of *P. incarnata* formulation was observed to be 2.5 mg quercetin equivalent /g of dry weight. The shaking test also showed the presence of saponin in the sample. The percentage discoloration of DPPH for the sample was found to be 87.25%, which was comparable to the % discoloration for ascorbic acid (93.56%). The total protein content in the *P. incarnata* sample was found to be 0.591 mg/mL. Qualitative tests also indicated the presence of saponin and carbohydrates in the *P. incarnata* formulation. Furthermore, LC-MS data of the *P. incarnata* formulation indicated the presence of the following compounds: 4-phenoxyphenol, Diphenhydramine, Cyclizine. It is evident that the *P. incarnata* formulation is rich in a variety of phytochemicals that may be responsible for altering sperm motility as discussed in the following sections.

### 3.4 *P. incarnata* formulation showed marked reduction in the sperm horizontal and vertical motility of sperm cells extracted from goat caudal epididymis

Goat sperm cells extracted from the caudal epididymis sections obtained from local slaughterhouses were used for the *ex vivo* studies. To check the effect of *P. incarnata* formulation on sperm motility, horizontal and vertical motility were studied using microscopic and spectrophotometric methods. A marked reduction in the horizontal motility was noted with the increase in the concentrations of the *P. incarnata* formulation from 40% to 11.5% (n = 3, *p* < 0.0001) (**Fig 2a**). A 1 mg/mL solution of aqueous *P. incarnata* formulation showed the optimum reduction in sperm horizontal motility and 0.1 mg/mL solution was optimum to reduce the vertical motility. This could be because of the combined action of the *P. incarnata* formulation and the action of gravity. There was a significant reduction in the % forward motility as the doses were increased gradually. Spectrophotometric analysis for measuring vertical motility revealed a significant decline in vertical motility after treatment with *P. incarnata* formulation (n = 3, *p* < 0.0001) (**Fig 2b**).

**Fig 2:**
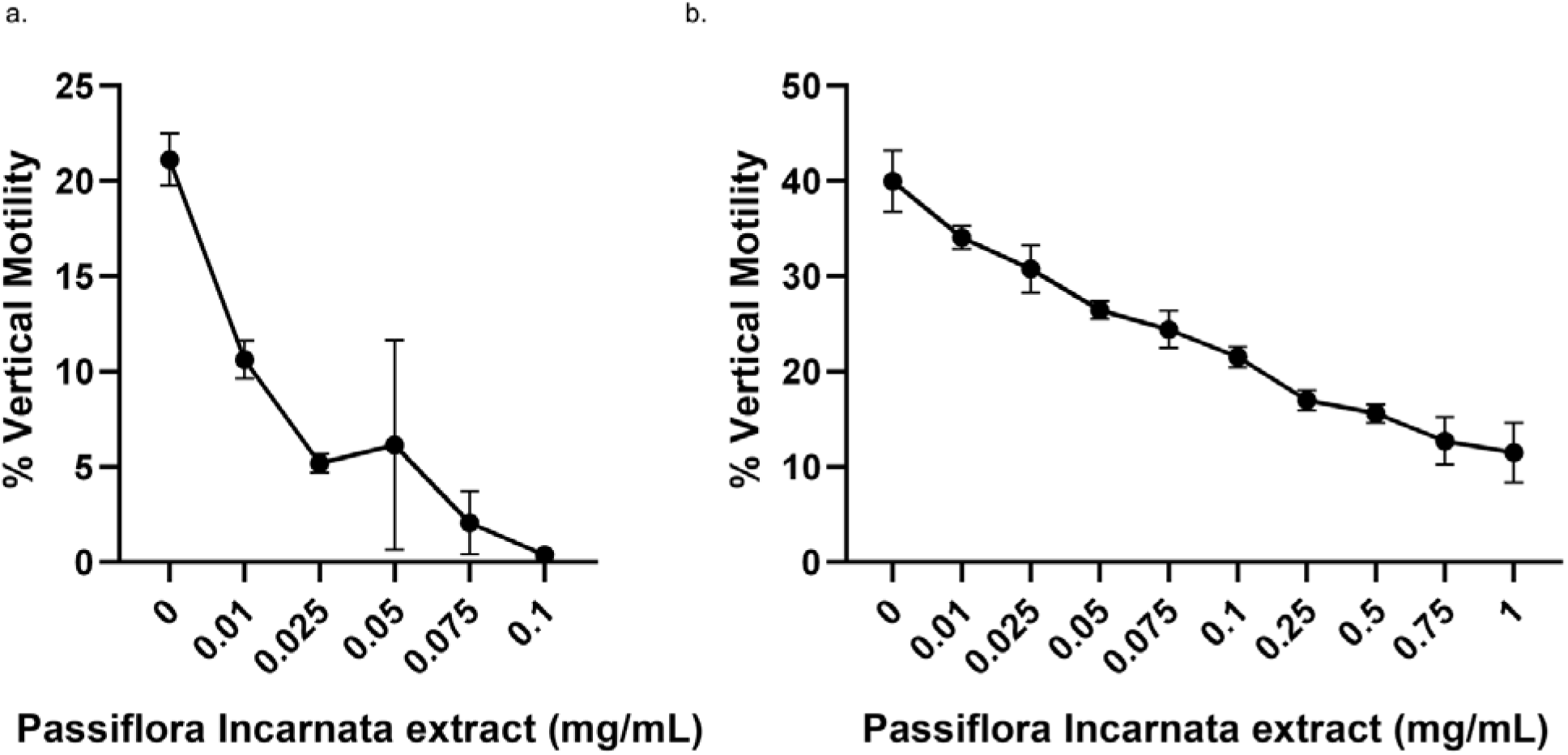
Effect of *P. incarnata* extract at different concentrations on sperm motility under the standard assay conditions. The values indicate the mean ± SEM of three experiments. A statistically significant difference was found with respect to the control (n = 3, *p* < 0.0001).

### 3.5 *P. incarnata* formulation did not affect the survivability of sperm cells; it inactivated motility selectively

Sperm viability was assessed using the trypan blue method (**Fig 3**). Interestingly, the results revealed no significant change in the viability of the cells, indicating that the *P. incarnata* formulation inhibited motility without killing the cells. There seems to be an underlying mechanism that is responsible for inhibiting sperm motility, without killing the cells. This mechanism needs to be assessed further.

**Fig 3:**
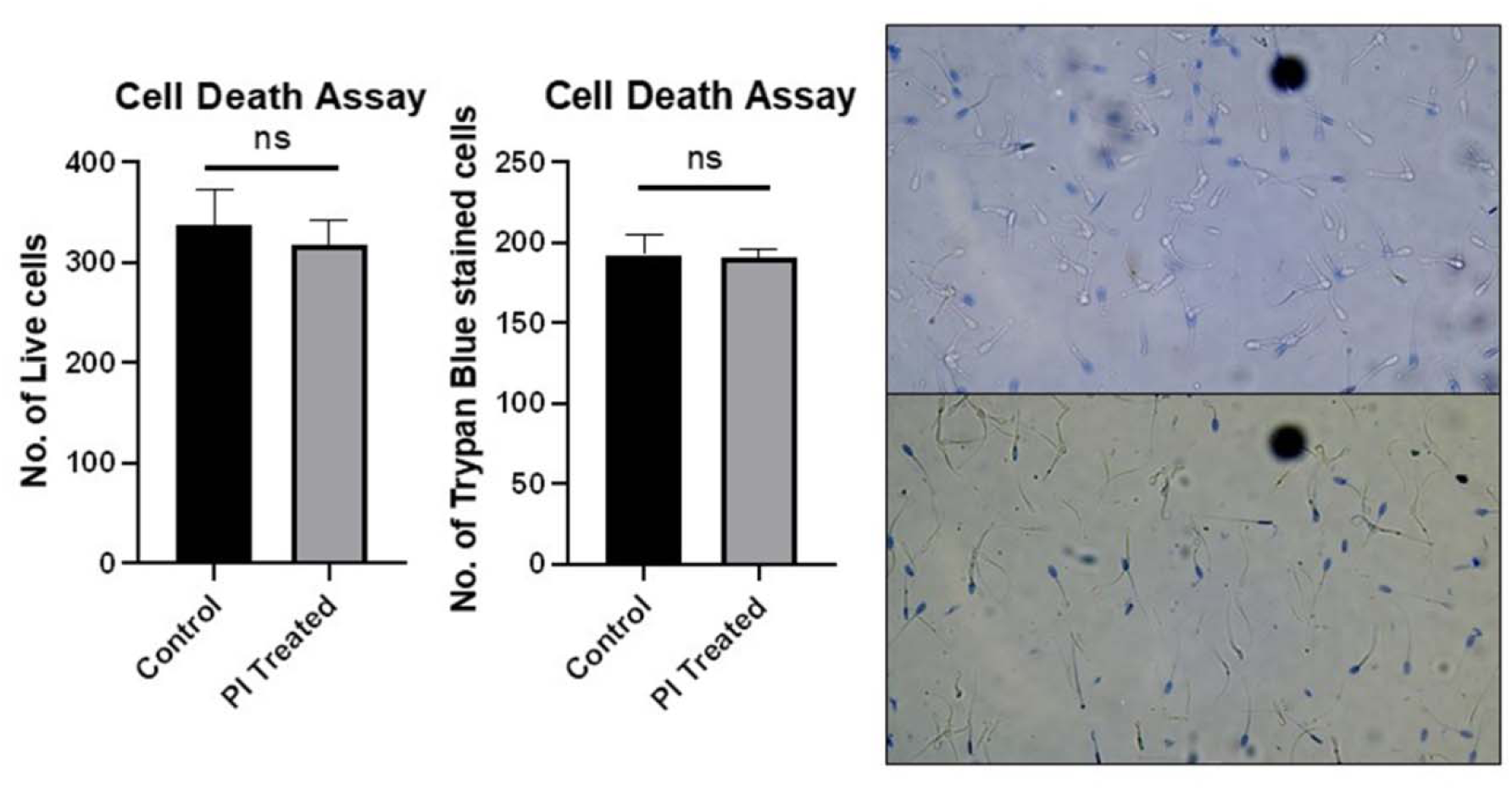
Trypan Blue staining of sperm cells after treatment with *Passiflora incarnata* extract showed no change in the number of stained cells (n = 3).

## 4. Discussion

It is known that most contraceptive methods are women-specific and generally target hormonal regulation [1]. So far very few studies have demonstrated the use of a male-centric, plant-based contraceptive. The observed dose-dependent inhibition of goat caudal epididymal sperm motility by *P. incarnata* formulation in our study aligns with emerging evidence on phytochemical modulation of sperm kinematics. We noted that horizontal motility declined from 40% to 11.5% (p < 0.0001) as *P. incarnata* formulation concentrations increased, while vertical motility exhibited greater sensitivity, requiring tenfold lower concentrations (0.1 mg/mL vs. 1 mg/mL) for optimal suppression. This disparity likely arises from gravitational strain during vertical ascent, which amplifies metabolic demands on spermatozoa. Spectrophotometric vertical velocity analysis corroborates findings from a previously described vertical motility analyzer, which quantifies motility against gravity through absorbance decay. Such methods overcome limitations of traditional horizontal motility assessments, as vertical movement better reflects *in vivo* challenges like cervical mucus penetration [17]. The cauda epididymidis microenvironment naturally suppresses motility via proteinaceous inhibitors like MIF-II (160 kDa), which lowers intracellular cAMP levels to maintain sperm quiescence [19]. *PI*’s phytochemicals-notably 16.85 mg GAE/g phenolics and saponins-likely exacerbate this suppression through dual mechanisms. Polyphenols such as quercetin analogs may disrupt redox-sensitive cAMP pathways, as demonstrated by their inhibition of human sperm tyrosine phosphorylation and calcium signaling [20]. Concurrently, saponins sequester membrane cholesterol, destabilizing lipid rafts essential for calcium influx and hyperactivation [21]. This synergistic action mirrors the motility-inhibiting effects of *Symplocos racemosa* saponins, which reduce membrane fluidity and ATPase activity in caprine models[22]. Moreover, the stark decline in forward motility (% reduction: p < 0.0001) parallels quercetin’s dose-responsive spermiostatic effects, where 100–400 μM concentrations inhibit human sperm motility via [Ca^2^□]□ depletion [20]. Our findings revealed a significant reduction in sperm motility following exposure to *Passiflora incarnata* formulation, shedding light on the potential contraceptive properties of this plant extract. *Passiflora incarnata* has been traditionally used for its anxiolytic and sedative properties, attributed to its diverse array of bioactive compounds [11]. While the impact on anxiety and sleep disorders has been widely studied, our research extends the understanding of the physiological effects of *Passiflora incarnata* to include reproductive processes. The biochemical composition of *Passiflora incarnata* formulation, as determined in our study, provides valuable insights into the potential mechanisms contributing to reduced sperm motility. *Passiflora incarnata* contains a variety of bioactive compounds, primarily flavonoids, alkaloids, phenols and phenolic acids. These compounds are linked to antioxidant activities. At high concentrations flavonoids such as chrysin may generate enough ROS that may peroxidize sperm membrane lipids and degrade flagellar proteins.

As per our findings we see that the aqueous formulation of *P. incarnata* demonstrated significant phytochemical diversity, with a total phenolic content of 16.85 mg gallic acid equivalents (GAE)/g dry weight and total flavonoid content of 2.5 mg quercetin equivalents (QE)/g dry weight. Saponins were qualitatively confirmed via foam testing, while DPPH radical scavenging assays revealed 87.25% antioxidant activity, comparable to ascorbic acid (93.56%). Liquid chromatography-mass spectrometry (LC-MS) identified 4-phenoxyphenol, diphenhydramine, and cyclizine as key constituents. These findings align with studies linking polyphenols and saponins to antioxidant and spermiomodulatory effects, suggesting *P. incarnata* formulation may influence sperm motility through redox regulation and bioactive compound interactions.

The Folin-Ciocalteu assay revealed a total phenolic content (TPC) of 16.85 ± 0.72 mg GAE/g dry weight in the *P. incarnata* formulation, consistent with methodologies validated for plant-derived matrices. Phenolic compounds, including flavonoids, act as primary antioxidants by donating hydrogen atoms to stabilize free radicals. The aluminum chloride colorimetric method quantified flavonoids at 2.5 ± 0.15 mg QE/g dry weight, a range observed in Musa paradisiaca tepal extracts (0.164–0.25 mg QE/g) [23,24]. Structural features such as hydroxyl group positioning in flavonoids enhance electron donation, critical for neutralizing reactive oxygen species (ROS). Phenolic compounds, a diverse class of phytochemicals, exert inhibitory effects on sperm motility through multifaceted mechanisms involving oxidative stress, membrane disruption, ion channel modulation, and metabolic interference. These interactions compromise sperm structural integrity, energy production, and signaling pathways essential for progressive movement. High phenolic concentrations disrupt redox homeostasis by increasing ROS production in spermatozoa. Compounds like quercetin and caffeic acid stimulate mitochondrial ROS generation via electron leakage from Complex I and III of the electron transport chain [25,26]. Simultaneously, phenolic metabolites such as quinones undergo redox cycling, producing superoxide radicals (O□□) and hydrogen peroxide (H□O□) [23]. This oxidative surge overwhelms endogenous antioxidants (e.g., glutathione, superoxide dismutase), leading to lipid peroxidation.

Saponins, a class of amphiphilic glycosides found in various plants, exhibit diverse biological activities, including spermicidal effects. Multiple studies demonstrate their capacity to impair sperm motility through membrane disruption, oxidative stress induction, and interference with critical signaling pathways. A primary mechanism of saponin-induced sperm immobilization involves their interaction with cholesterol in the sperm plasma membrane. Saponins such as digitonin form complexes with membrane cholesterol, destabilizing lipid bilayers and creating pores that compromise membrane integrity [27]. The loss of membrane integrity disrupts ion homeostasis, particularly calcium (Ca^2^□) flux, which is critical for hyperactivation and chemotaxis. By increasing membrane permeability, saponins induce uncontrolled Ca^2^ influx, leading to premature acrosome reactions and impaired motility [28].

Plant-derived flavonoids exhibit complex, concentration-dependent effects on sperm motility, ranging from protective antioxidant activity at low doses to inhibitory or spermicidal actions at higher concentrations. The *P. incarnata* formulation exhibited 87.25% DPPH radical inhibition at 50 μg/mL, nearing ascorbic acid’s 93.56%. This dose-dependent scavenging aligns with *Cordia dichotoma* methanol extracts (IC_50_ = 28 μg/mL) [29]. The DPPH assay’s mechanism involves electron transfer from antioxidants to the stable radical, forming a reduced, pale-yellow product. While antioxidants are generally associated with beneficial effects on cellular health, their role in sperm motility is complex. Some studies suggest that moderate levels of oxidative stress are necessary for normal sperm function, and an excess of antioxidants may disrupt this delicate balance. [1,14,30].

Flavonoids directly compromise sperm membrane integrity through two primary mechanisms. Amphiphilic flavonoids like ursolic acid (a triterpenoid) bind membrane cholesterol, forming pore-like structures that increase permeability. This disrupts lipid raft organization critical for calcium signalling and acrosome reaction initiation. Transmission electron microscopy reveals flavonoid-treated sperm exhibit acrosomal swelling and plasma membrane fragmentation [14]. Pro-oxidant flavonoids (e.g., quercetin) at high concentrations (>50 μM) paradoxically induce reactive oxygen species (ROS) that peroxidize sperm membrane polyunsaturated fatty acids (PUFAs). Malondialdehyde (MDA) levels rise 3-fold in human sperm exposed to *Ziziphus mauritiana* flavonoids, correlating with complete motility loss at 0.5 mg/mL [31]. High flavonoid concentrations directly damage motility structures and cause axonemal dysfunction. A study shows that quercetin (100 μM) cross-links tubulin dimers and reduces microtubule sliding velocity by 70% in demembranated sperm models [32]. Flavonoids also cause fibrous sheath disintegration. A study indicated that Catechin (500 μM) degrades AKAP3 scaffolding proteins, disrupting protein kinase A (PKA) localization critical for flagellar bending [33].

4-Phenoxyphenol (PhOP), a para-alkoxyphenol, may directly impair sperm function through oxidative stress. Phenolic compounds often generate reactive oxygen species (ROS), which peroxidize sperm membrane lipids, deplete antioxidants like glutathione (GSH), and damage mitochondrial ATP production . While specific data on PhOP’s pro-oxidant effects are limited, structurally similar polyphenols induce superoxide radicals that immobilize sperm by degrading flagellar proteins and disrupting ion channels [23,34]. Diphenhydramine is a first-generation H1 antihistamine and may inhibit sperm motility by regulating the calcium signaling. Histamines regulate sperm calcium (Ca^2^□) dynamics through H1 receptors on the sperm membrane. Binding of histamine to H1 receptors activates phospholipase C (PLC), generating inositol trisphosphate (IP□), which triggers Ca^2^□ release from intracellular stores. Diphenhydramine antagonizes H1 receptors, preventing histamine-induced Ca^2^□ elevation. At low doses (0.01–1.0 mM), this blockade protects against histamine’s spermicidal effects but disrupts physiological Ca^2^□ oscillations required for hyperactivation and acrosomal exocytosis [35]. At higher concentrations (>10 mM), diphenhydramine becomes spermicidal, inducing membrane lipid peroxidation and irreversible Ca^2^□ overload. This disrupts mitochondrial membrane potential, halting ATP production and paralyzing flagellar movement. In hypokinetic human sperm, diphenhydramine reduces viability by 80% within 60 minutes, correlating with malondialdehyde (MDA) accumulation [36,37]. Cyclizine, another H1 antagonist, exerts cytotoxicity through apoptosis induction in mammalian cells. In RAW264.7 macrophages, cyclizine activates both intrinsic and extrinsic apoptotic pathways through mitochondrial depolarization wherein there is a reduction in the Bcl-2/Bad protein ratio, triggering cytochrome C release. Cyclizine also causes an upregulation in caspases-3, −8, and −9. Furthermore, it may also modulate death receptor thereby promoting the levels of Enhancing tumor necrosis factor-α receptor (TNFR) and Fas receptor expression, promoting sperm cell death [38]. Similarly, in sperm, these pathways likely disrupt Ca^2^ homeostasis and energy metabolism, immobilizing cells via ATP depletion and cytoskeletal degradation, specifically structural proteins like dynein arms in the sperm flagellum. Cyclizine’s adduct profile in LC-MS (e.g., m/z 167.01 fragment) further suggests interactions with motility-related proteins.

The flavonoids (chrysin, BZF, vitexin) and phenolic acids in *Passiflora incarnata* primarily influence sperm motility via antioxidant pathways. Alkaloids like harmine may contribute indirectly through genotoxic effects. Dose-dependent biphasic effects underscore the need for further research to optimize therapeutic applications while minimizing risks. While our study provides valuable insights into the potential contraceptive properties of *Passiflora incarnata*, it is essential to acknowledge certain limitations. The complexity of the biochemical composition of the formulation makes it challenging to pinpoint specific molecules responsible for the observed effects. Future research should focus on isolating individual compounds from this formulation and assessing their impact on sperm motility to unravel the precise mechanisms at play.

## 5. Conclusion

This study highlights the complex biological macromolecules such as polyphenols in *Passiflora incarnata* aqueous formulation and its ability to significantly reduces sperm motility, suggesting its potential as a natural contraceptive agent. The high phenolic, saponin, flavonoid, and protein content of the formulation provides a biochemical basis for these observed effects. Being water soluble, lubricant gel may be prepared in future which may be applied during intercourse as well for the couple who wish to prevent unplanned pregnancy. This research contributes to the growing body of knowledge on the contraceptive potential of plant extracts and opens avenues for further exploration into the development of botanical-based contraceptives.

## Acknowledgment

The authors would like to thank and acknowledge the analytical and technical support by Dr. Arpita Bhoumik, CSIR-Indian Institute of Chemical Biology, Kolkata, India. The authors are thankful to the Amity Institute of Physiology and Allied Sciences and Amity Institute of Pharmacy, Amity University Uttar Pradesh, India for providing necessary infrastructure and facilities to perform the experiments. The authors would also like to acknowledge the Department of Biotechnology, Sister Nivedita University, Kolkata and the kind support of the Vice Chancellor, Sister Nivedita University, Kolkata, India.

## Funding

This work was supported by the Science and Engineering Research Board Start-up Research Grant [grant number: SRG/2019/000501]

## Statement of declarations Competing interests

The authors declare no conflict of interests.

## Authors’ contributions

SC contributed to the investigation, methodology, resource acquisition, data interpretation, and writing of the original draft. HKS was involved in investigation and methodology. Both SC and HKS are joint first authors of the article. SS was responsible for project administration, resource management, funding acquisition. KN and SS led the conceptualization and formal analysis, participated in the investigation, provided supervision and visualization support, and were responsible for writing, reviewing and editing the article as well as serving as the joint corresponding authors.

## Availability of data and material

Not applicable.

## Clinical trial number

Not applicable

## Ethics, Consent to Participate, and Consent to Publish declarations

Not applicable.

## Notes

### Competing Interest Statement

The authors have declared no competing interest.

## References

1. Chakraborty S, Saha S. Understanding sperm motility mechanisms and the implication of sperm surface molecules in promoting motility. Middle East Fertil Soc J. 2022;27:4.

2. Louwagie EJ, Quinn GFL, Pond KL, Hansen KA. Male contraception: narrative review of ongoing research. Basic Clin Androl. 2023;33:30.

3. Hifnawy MS, Aboseada MA, Hassan HM, Tohamy AF, El Naggar EMB, Abdelmohsen UR. Nature-inspired male contraceptive and spermicidal products. Phytochem Rev. 2021;20:797–843.

4. Teal S, Edelman A. Contraception Selection, Effectiveness, and Adverse Effects: A Review. JAMA. 2021;326:2507–18.

5. Williamson LM, Parkes A, Wight D, Petticrew M, Hart GJ. Limits to modern contraceptive use among young women in developing countries: a systematic review of qualitative research. Reprod Health. 2009;6:3.

6. Rosenberg MJ, Waugh MS, Meehan TE. Use and misuse of oral contraceptives: Risk indicators for poor pill taking and discontinuation. Contraception. 1995;51:283–8.

7. Kumar D, Kumar A, Prakash O. Potential antifertility agents from plants: A comprehensive review. J Ethnopharmacol. 2012;140:1–32.

8. Moroole MA, Materechera SA, Otang-Mbeng W, Hayeshi R, Bester C, Aremu AO. Phytochemical Profile, Safety and Efficacy of a Herbal Mixture Used for Contraception by Traditional Health Practitioners in Ngaka Modiri Molema District Municipality, South Africa. Plants. 2022;11:193.

9. Ogbuewu IP, Unamba-Oparah IC, Odoemenam VU, Etuk IF, Okoli IC. The potentiality of medicinal plants as the source of new contraceptive principles in males. North Am J Med Sci. 2011;3:255–63.

10. Janda K, Wojtkowska K, Jakubczyk K, Antoniewicz J, Skonieczna-Żydecka K. Passiflora incarnata in Neuropsychiatric Disorders—A Systematic Review. Nutrients. 2020;12:3894.

11. Kim G-H, Yi SS. Chronic oral administration of Passiflora incarnata extract has no abnormal effects on metabolic and behavioral parameters in mice, except to induce sleep. Lab Anim Res. 2019;35:31.

12. Guerrero FA, Medina GM. Effect of a medicinal plant (Passiflora incarnata L) on sleep. Sleep Sci. 2017;10:96–100.

13. Singh HK, Nagpal DK. The UV spectrophotometric based analytical method development and validation for the quantitative estimation of Passiflora incarnata. SGS - Eng Sci [Internet]. 2021 [cited 2025 Apr 8];1. Available from: https://spast.org/techrep/article/view/2643

14. Chattopadhyay D, Dungdung SR, Mandal AB, Majumder GC. A potent sperm motility-inhibiting activity of bioflavonoids from an ethnomedicine of Onge, Alstonia macrophylla Wall ex A. DC, leaf extract. Contraception. 2005;71:372–8.

15. Saha S, Das S, Bhoumik A, Ghosh P, Majumder GC, Dungdung SR. Identification of a novel sperm motility–stimulating protein from caprine serum: its characterization and functional significance. Fertil Steril. 2013;100:269–279.e5.

16. Roy N, Majumder GC, Chakrabarti CK. Occurrence of Specific Glycoprotein Factor(s) in Goat Epididymal Plasma that Prevent Adhesion of Spermatozoa to Glass. Andrologia. 1985;17:200–6.

17. Saha S, Paul D, Mukherjee A, Banerjee S, Majumder GC. A computerized spectrophotometric instrumental system to determine the “vertical velocity” of sperm cells: A novel concept. Cytometry A. 2007;71A:308–16.

18. Mohammadi A, Kanfer I, Sewram V, Walker RB. An LC-MS-MS method for the determination of cyclizine in human serum. J Chromatogr B Analyt Technol Biomed Life Sci. 2005;824:148–52.

19. Das S, Saha S, Majumder GC, Dungdung SR. Purification and characterization of a sperm motility inhibiting factor from caprine epididymal plasma. PloS One. 2010;5:e12039.

20. Liang X, Xia Z, Yan J, Wang Y, Xue S, Zhang X. Quercetin inhibits human sperm functions by reducing sperm [Ca2+]i and tyrosine phosphorylation. Pak J Pharm Sci. 2016;29:2391–6.

21. Vinarova L, Vinarov Z, Atanasov V, Pantcheva I, Tcholakova S, Denkov N, et al. Lowering of cholesterol bioaccessibility and serum concentrations by saponins: in vitro and in vivo studies. Food Funct. 2015;6:501–12.

22. Lal P, Jorasia K, Rathore NS, Kumar V, Singh R, Moolchandrani A, et al. Purification and partial characterization of a sperm motility inhibitory protein of ram cauda epididymal plasma. Cell Biochem Funct. 2024;42:e3930.

23. Kumar P, Laloraya M, Laloraya MM. The effect of some of the polyphenolic compounds on sperm motility in vitro: a structure-activity relationship. Contraception. 1989;39:531–9.

24. Pérez M, Dominguez-López I, Lamuela-Raventós RM. The Chemistry Behind the Folin-Ciocalteu Method for the Estimation of (Poly)phenol Content in Food: Total Phenolic Intake in a Mediterranean Dietary Pattern. J Agric Food Chem. 2023;71:17543–53.

25. Aitken RJ, Muscio L, Whiting S, Connaughton HS, Fraser BA, Nixon B, et al. Analysis of the effects of polyphenols on human spermatozoa reveals unexpected impacts on mitochondrial membrane potential, oxidative stress and DNA integrity; implications for assisted reproductive technology. Biochem Pharmacol. 2016;121:78–96.

26. Castellini C, D’Andrea S, Cordeschi G, Totaro M, Parisi A, Di Emidio G, et al. Pathophysiology of Mitochondrial Dysfunction in Human Spermatozoa: Focus on Energetic Metabolism, Oxidative Stress and Apoptosis. Antioxid Basel Switz. 2021;10:695.

27. Sudji I, Subburaj Y, Frenkel N, García-Sáez A, Wink M. Membrane Disintegration Caused by the Steroid Saponin Digitonin Is Related to the Presence of Cholesterol. Molecules. 2015;20:20146–60.

28. Rahman MS, Kwon W-S, Pang M-G. Calcium influx and male fertility in the context of the sperm proteome: an update. BioMed Res Int. 2014;2014:841615.

29. Nariya PB, Bhalodia NR, Shukla VJ, Acharya R, Nariya MB. In vitro evaluation of antioxidant activity of Cordia dichotoma (Forst f.) bark. Ayu. 2013;34:124–8.

30. Nass-Arden L, Breitbart H. Modulation of mammalian sperm motility by quercetin. Mol Reprod Dev. 1990;25:369–73.

31. Kaltsas A. Oxidative Stress and Male Infertility: The Protective Role of Antioxidants. Med Kaunas Lith. 2023;59:1769.

32. Jamalan M, Ghaffari MA, Hoseinzadeh P, Hashemitabar M, Zeinali M. Human Sperm Quality and Metal Toxicants: Protective Effects of some Flavonoids on Male Reproductive Function. Int J Fertil Steril [Internet]. 2016 [cited 2025 May 14];10. Available from: 10.22074/ijfs.2016.4912

33. Purdy PH, Ericsson SA, Dodson RE, Sternes KL, Garner DL. Effects of the flavonoids, silibinin and catechin, on the motility of extended cooled caprine sperm. Small Rumin Res. 2004;55:239–43.

34. Schiffer C, Müller A, Egeberg DL, Alvarez L, Brenker C, Rehfeld A, et al. Direct action of endocrine disrupting chemicals on human sperm. EMBO Rep. 2014;15:758–65.

35. Gupta A, Khosla R, Gupta S, Tiwary AK. Influence of histamine and H1-receptor antagonists on ejaculated human spermatozoa: role of intrasperm Ca2+. Indian J Exp Biol. 2004;42:481–5.

36. Adamczak R, Ukleja-Sokołowska N, Pasińska M, Zielińska J, Leśny M, Dubiel M. Abnormal sperm morphology is associated with sensitization to inhaled allergens. Int J Immunopathol Pharmacol. 2022;36:20587384211066718.

37. Mondillo C, Varela ML, Abiuso AMB, Vázquez R. Potential negative effects of anti-histamines on male reproductive function. Reprod Camb Engl. 2018;155:R221–7.

38. Lu Y, Chiang C, Hsu Y, Chen C, Chen W, Tseng C, et al. Cyclizine induces cytotoxicity and apoptosis in macrophages through the extrinsic and intrinsic apoptotic pathways. Environ Toxicol. 2024;39:2970–9.

